# Dissociable control of persistence and adaptive updating in reward-guided behavior

**DOI:** 10.64898/2026.09.04.749383

**Authors:** Pratheba Kandasamey, Anaïs Bouchat, Ladina Odermatt, Eva Bracey, Denis Burdakov, Daria Peleg-Raibstein

## Abstract

Adaptive behavior requires a balance between persistence and flexibility, whereas excessive persistence and impaired updating characterize several neuropsychiatric disorders. Touchscreen-based cognitive tasks provide a reverse-translational framework because closely related visual-discrimination and reversal paradigms are used to phenotype cognitive flexibility in rodents and humans. Here, we examined the role of the melanin-concentrating hormone (MCH) system using appetitive touchscreen extinction and reversal tasks. Blocking MCHR1 had little effect while an established reward contingency remained valid, but facilitated behavioral adaptation when reinforcement was withdrawn or reversed. These findings suggest that MCHR1 signaling promotes persistence of previously reinforced behavior. We then chemogenetically activated lateral hypothalamic MCH neurons. Activation produced no detectable change in extinction, but altered learning during reversal. Notably, behavioral differences were also evident during a subsequent reversal performed without chemogenetic activation: mice previously exposed to MCH-neuron activation learned the new contingency more rapidly, yet showed greater persistence after individual errors. Thus, MCHR1 blockade and MCH-neuron activation affect distinct components of behavioral updating rather than producing simple opposite effects. Together, these findings identify dissociable MCH-system contributions to reward-contingency updating in a behavioral framework directly relevant to cross-species neuropsychiatric research.

## Introduction

Adaptive behavior requires learned associations to remain sufficiently stable to guide action while retaining the capacity to be revised when their consequences change. This balance becomes particularly important when an established response is no longer adaptive. During extinction, omission of an expected outcome progressively reduces expression of a previously reinforced response, whereas reversal learning requires suppression of the previously valid response together with acquisition of a competing contingency. Reversal therefore places additional demands on behavioral control because past and current reinforcement histories compete to determine choice (Cools et al., 2002; Fellows and Farah, 2003; Remijnse et al., 2005; Ragozzino, 2007; Floresco et al., 2009; Bissonette and Powell, 2012). Failures of this balance are prominent in neuropsychiatric conditions characterized by compulsive or perseverative behavior, including obsessive–compulsive disorder and addiction (Remijnse et al., 2006; Izquierdo and Jentsch, 2012).

Computerized reversal-learning tasks are used in humans to probe cognitive flexibility and reveal abnormalities in neuropsychiatric populations (Cools et al., 2002; Remijnse et al., 2005, 2006). Rodent touchscreen tasks provide a reverse-translational counterpart, preserving comparable visual stimulus presentation, direct response selection and automated task structure across species (Bussey et al., 2012; Horner et al., 2013; Turner et al., 2017; Sullivan et al., 2021; Lopez-Cruz et al., 2021). Closely matched, and in some cases identical, tasks have been applied in rodents and humans, linking circuit-level studies to clinical cognitive phenotyping (Nithianantharajah et al., 2015; Palmer et al., 2021). Visual discrimination and reversal within this platform therefore allow neurobiological mechanisms of contingency updating to be examined in a format with direct translational relevance.

Melanin-concentrating hormone (MCH) neurons, located in the lateral hypothalamic area and adjacent zona incerta, form a widely projecting system classically associated with energy balance, feeding, sleep and reward (Bittencourt et al., 1992; Qu et al., 1996; Shimada et al., 1998; Georgescu et al., 2005; Hassani et al., 2009; Diniz and Bittencourt, 2019; Al-Massadi et al., 2021; Lord et al., 2021). A broader literature now implicates MCH neurons in experience-dependent control of behavior. MCH-neuron activity is recruited during appetitive and consummatory behavior (Subramanian et al., 2023; Potter et al., 2026), while MCH signaling has been linked to object memorization, hippocampus-dependent memory persistence, fear extinction and response inhibition (Kosse and Burdakov, 2019; Izawa et al., 2019; Noble et al., 2019; Concetti et al., 2020). Together, these findings suggest that the MCH system may influence not only reward-related behavior, but also the persistence and modification of previously acquired information (Burdakov and Peleg-Raibstein, 2020; Burdakov and Peleg-Raibstein, 2025; Concetti et al., 2024).

An important distinction, however, is that chemogenetic activation of MCH neurons and signaling through MCHR1 are not functionally equivalent manipulations. MCH neurons engage multiple neurochemical outputs, including glutamatergic and peptidergic signaling with dissociable physiological effects (Schneeberger et al., 2018), whereas MCHR1 antagonism selectively interferes with receptor-mediated MCH signaling. Consistent with this distinction, increasing MCH-neuron activity and chronically reducing MCH/MCHR1 signaling can produce related rather than simply opposing behavioral consequences (Noble et al., 2019). Thus, the contribution of MCHR1 signaling to behavioral updating cannot be inferred directly from the consequences of manipulating MCH-neuron activity.

Recent work suggests that MCHR1 signaling supports the persistence of previously reinforced behavior when contingencies change. In active avoidance, MCHR1 antagonism facilitated extinction of a learned defensive response, while in a reward-guided T-maze it reduced perseverative responding and accelerated adaptation following contingency reversal (Kandasamey et al., 2026a,b). Together, these findings support a role for MCHR1 signaling in the updating of established behavior across aversive and appetitive contexts.

Here, we asked whether MCH-system control of reward-guided behavior emerges preferentially when established contingencies must be revised, and whether receptor-level MCHR1 signaling and chemogenetic activation of LH-MCH neurons contribute to this process in the same way. By comparing extinction, in which reinforcement is withdrawn, with reversal, in which an obsolete contingency must be replaced by a competing rewarded alternative, we sought to define how distinct levels of the MCH system regulate the balance between behavioral persistence and adaptive updating.

## Results

### MCHR1 antagonism most strongly facilitates extinction when present during contingency change

Our previous work implicated MCHR1 signaling in the persistence of established behavior across aversive and appetitive settings (Kandasamey et al., 2026a,b). We therefore asked whether this contribution depends on when receptor signaling is disrupted by administering the selective MCHR1 antagonist SNAP-94847 during either acquisition or extinction of an appetitive touchscreen response (Fig. 1A,B). During acquisition, a transient visual stimulus was presented in a touchscreen response window and touching the stimulus was rewarded; during extinction, the same response was no longer reinforced. Control mice received vehicle during both phases, SNAP-Acq mice received SNAP during acquisition only, and SNAP-Ext mice received SNAP during extinction only.

**Figure 1.**
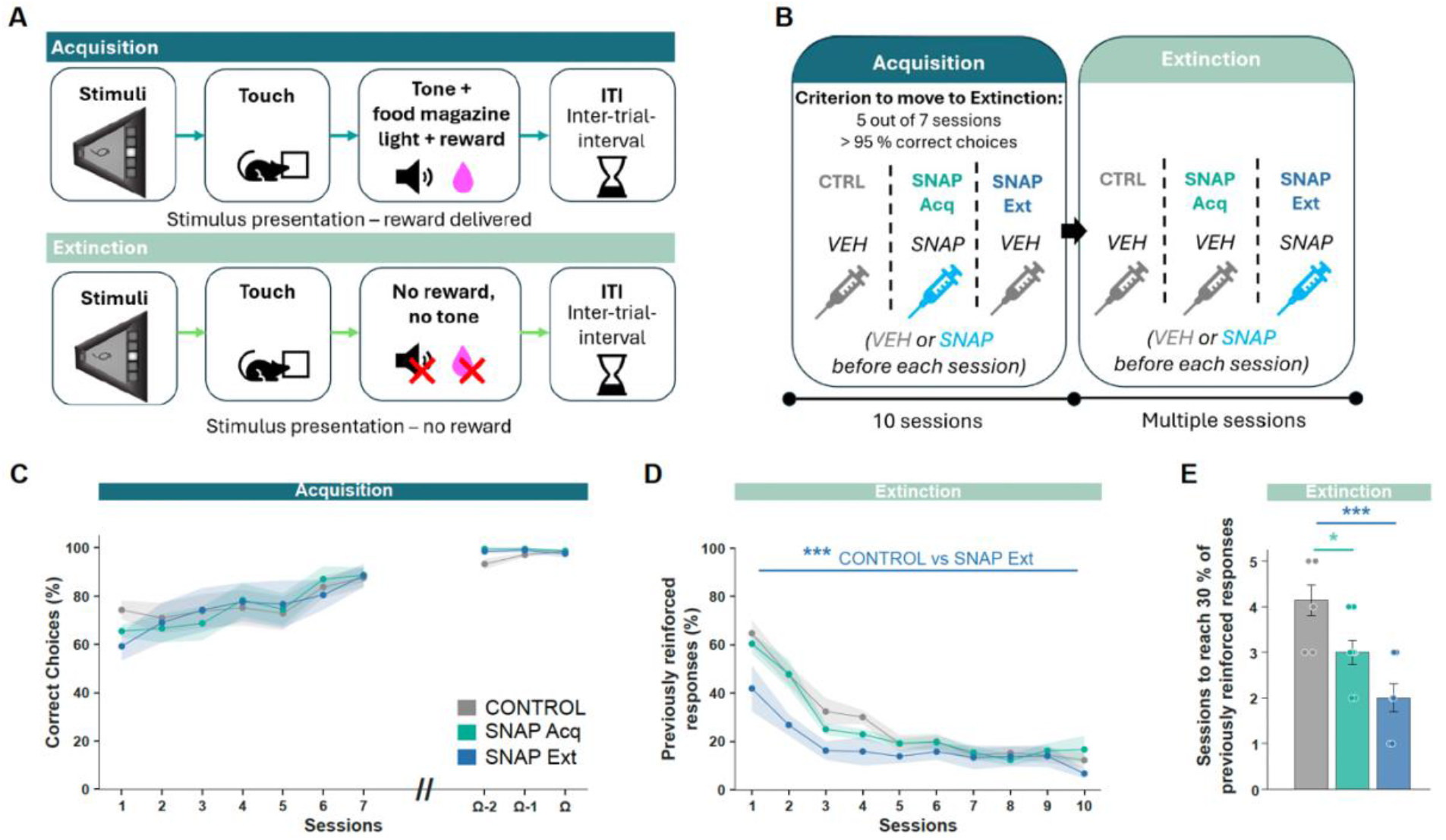
MCHR1 antagonism most strongly facilitates extinction when present during contingency change. (A) Touchscreen acquisition and extinction task; icons indicate the auditory cue (speaker), magazine light (bulb) and strawberry milkshake reward (pink droplet). (B) Experimental design. Mice received vehicle throughout (CONTROL), SNAP-94847 during acquisition only (SNAP Acq), or SNAP-94847 during extinction only (SNAP Ext). (C) Acquisition performance; Ω denotes the individual criterion session; Ω−1 and Ω−2 denote the one and two preceding sessions, respectively. (D) Responding to the previously reinforced stimulus during the first 10 extinction sessions. (E) Sessions required to reach ≤30% previously reinforced responding. Session trajectories were analyzed with linear mixed-effects models; criterion comparisons used Welch’s *t*-tests with FDR correction. Lines and shading show mean ± SEM; bars show mean ± SEM with individual animals. *p < 0.05, **p < 0.01, ***p < 0.001, ****p < 0.0001. (n CONTROL = 7, n SNAP Acq = 8, n SNAP Ext = 7)

Acquisition performance increased across sessions and did not differ among groups (session: β = 2.452, p < 0.001; treatment and treatment × session, all p > 0.40; Fig. 1C), indicating that SNAP administered during acquisition did not measurably impair establishment of the reinforced response. The groups diverged once reward was withdrawn. SNAP administered during extinction markedly reduced responding to the previously reinforced stimulus and altered the extinction trajectory relative to controls (group: β = −23.388, p < 0.001; group × session: β = 2.612, p < 0.001; Fig. 1D). SNAP administered during acquisition only did not produce a comparable trajectory effect (group: β = −4.977, p = 0.281; group × session: β = 0.543, p = 0.466), and the combined model confirmed an extinction-specific effect of SNAP-Ext (phase × SNAP-Ext × session: β = 2.898, p < 0.001).

Criterion attainment revealed a smaller carry-over effect of acquisition treatment: both SNAP-Acq and SNAP-Ext mice reached ≤30% previously reinforced responding earlier than controls (SNAP-Acq, pFDR = 0.0217; SNAP-Ext, pFDR = 0.00112; Fig. 1E). Thus, acquisition treatment was not without later consequence, but the dominant effect on extinction dynamics occurred when MCHR1 blockade coincided with the change in reinforcement contingency.

We next asked whether MCHR1 blockade also facilitated adaptation when the reward contingency was reversed.

### MCHR1 antagonism facilitates adaptation to reversal without reducing post-error perseveration

Having found that MCHR1 blockade was most effective when it coincided with extinction, we next tested its effect on reversal learning, in which the previously rewarded stimulus becomes unrewarded and its alternative becomes rewarded. Mice first acquired the visual discrimination to criterion before any manipulation of reversal. On each trial, two visual stimuli were presented in the touchscreen response windows, with one stimulus (S+) rewarded and the other (S−) unrewarded irrespective of screen location; during reversal, these stimulus–reward assignments were switched. Before reversal began, SNAP or vehicle was administered during one additional session in which the established discrimination remained unchanged. This pre-reversal challenge served as a control for acute treatment effects under a stable contingency: neither first-choice performance (accuracy on regular trials, excluding correction trials; between-group change: Welch’s t(10.58) = −1.22, pFDR = 0.374) nor correction-trial burden (Welch’s t(8.12) = 0.07, pFDR = 0.947) changed differentially between groups (Supplementary Fig. 1A,B). SNAP or vehicle was then administered before each reversal session (Fig. 2A,B).

**Figure 2.**
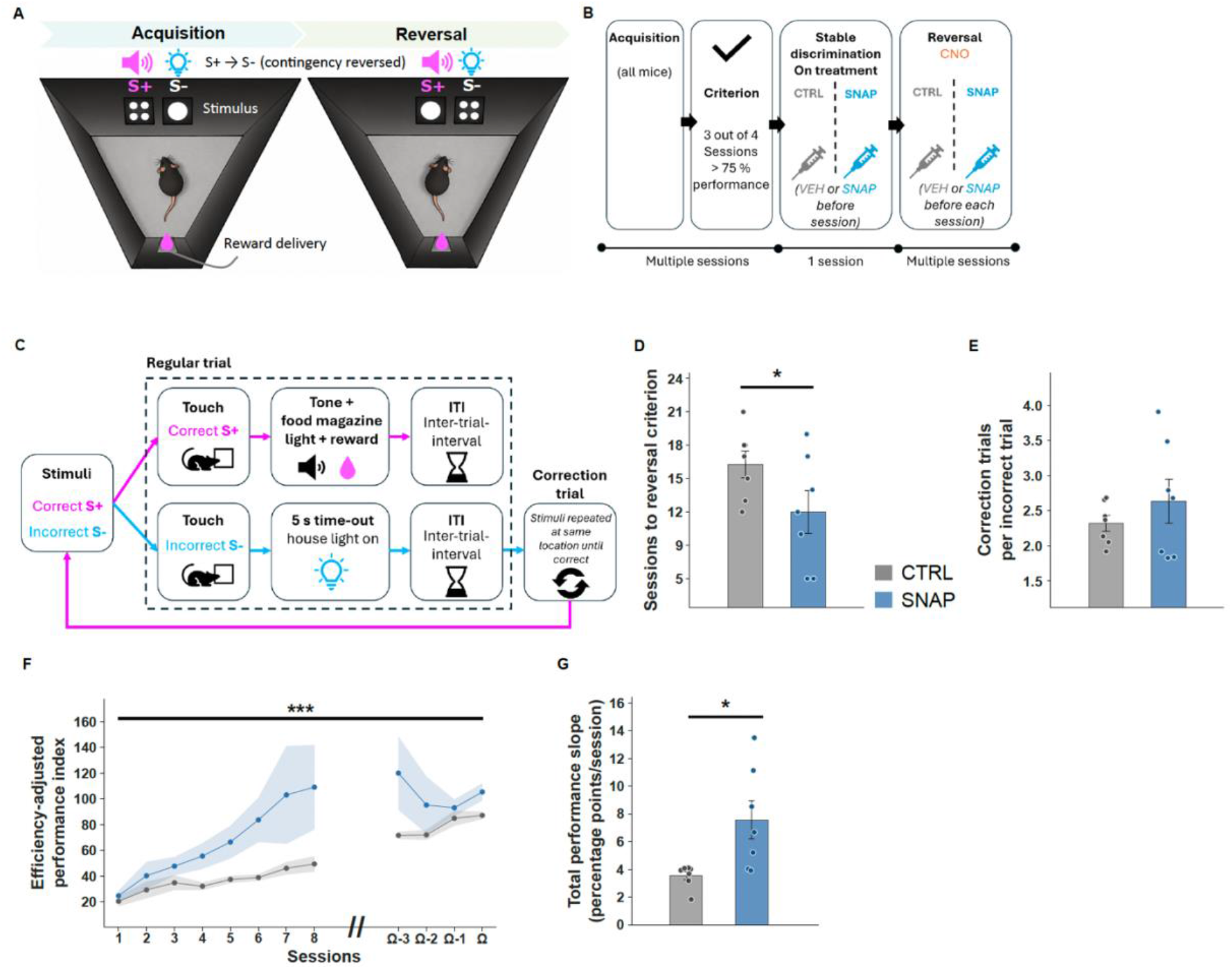
MCHR1 antagonism facilitates adaptation to reversal without reducing post-error perseveration. (A) Visual-discrimination acquisition and reversal task. S+, rewarded stimulus; S−, unrewarded stimulus. (B) Experimental design. Following acquisition to criterion, mice received SNAP or vehicle during one pre-reversal stable-discrimination challenge session, followed by the same treatment before each reversal session. (C) Correction-trial procedure. A correct regular-trial choice was rewarded and followed by the next trial, whereas an incorrect choice produced a time-out and repetition of the same stimulus configuration until a correct response was made. Correction trials were excluded from first-choice accuracy. (D) Sessions to reversal criterion. (E) Correction trials per incorrect regular trial. (F) Efficiency-adjusted performance index across reversal. For visualization, the first eight sessions and the final four criterion-aligned sessions are shown; all available reversal sessions were included in the cluster-robust OLS analysis. (G) Per-animal total-performance slope across reversal, estimated by linear regression of session-wise total performance, including regular and correction trials, across all available reversal sessions. Data are mean ± SEM. *p < 0.05, **p < 0.01, ***p < 0.001, ****p < 0.0001.

Once the contingency was reversed, SNAP-treated mice adapted more rapidly. They required fewer sessions to criterion (Control: 16.29 ± 1.19; SNAP: 12.00 ± 1.91; one-tailed Welch’s t(10.03) = 1.90, p = 0.043; a priori directional hypothesis; Fig. 2D). We next asked whether this faster criterion attainment reflected reduced persistence following an error. In touchscreen reversal, an incorrect regular trial was followed by repetition of the same stimulus configuration, including stimulus locations, until the correct stimulus was selected (Fig. 2C). Correction trials relative to initial errors therefore provide an established measure of post-error perseveration and response strategy (Turner et al., 2017; Marquardt et al., 2017; Dumont et al., 2025). Error-triggered behavioral adjustment likewise constitutes a separable component of cognitive control in humans (Rabbitt, 1966; Holroyd and Coles, 2002; Kerns et al., 2004; van Veen and Carter, 2006; Danielmeier and Ullsperger, 2011; Markovitch et al., 2024). SNAP did not alter correction trials per incorrect response (Control: 2.32 ± 0.11; SNAP: 2.64 ± 0.32; p = 0.373; Fig. 2E).

The faster criterion attainment was therefore not accompanied by a detectable reduction in post-error perseveration. Instead, SNAP-treated mice showed a higher efficiency-adjusted performance index, a trial-count-adjusted measure of total performance (group: β = 41.19, p < 0.001; Fig. 2F), and steeper per-animal total-performance slopes across reversal (Control: 3.55 ± 0.31; SNAP: 7.57 ± 1.39 percentage points/session; Welch’s t(6.60) = −2.83, p = 0.027; Fig. 2G). The efficiency-adjusted performance index showed a group difference without evidence for a group × session interaction (p = 0.176), consistent with a performance difference across reversal rather than a detectable change in its session dependence. Regular-trial accuracy itself did not differ between groups across reversal: first-choice performance increased with session, but neither the group effect nor the group × session interaction was significant (Supplementary Fig. 1C).

Having established a role for MCHR1 signaling in contingency updating, we next asked how chemogenetic activation of the MCH neuronal population influences the same behavioral process.

### Chemogenetic activation of LH-MCH neurons produces no detectable change in appetitive extinction

Having identified a preferential involvement of MCHR1 signaling when an established contingency was no longer reinforced, we next asked how chemogenetic activation of LH-MCH neurons influences the same behavioral transition. To isolate chemogenetic activation during contingency change from effects on formation of the original association, MCH-DREADD activation was restricted to extinction; acquisition was conducted without chemogenetic manipulation (Fig. 3A).

**Figure 3.**
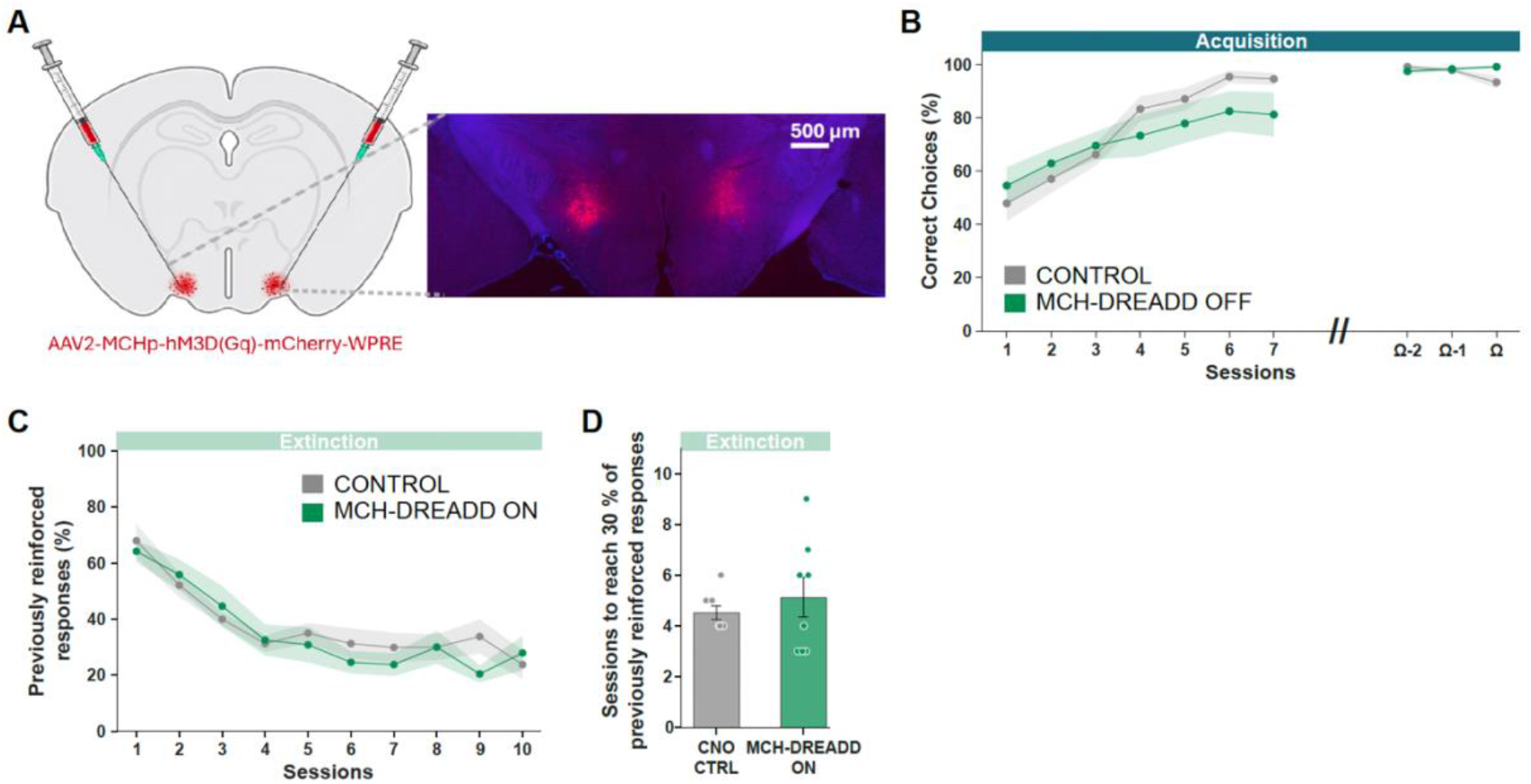
Chemogenetic activation of LH-MCH neurons produces no detectable change in appetitive extinction. (A) Viral targeting strategy and representative hM3Dq-mCherry expression in the lateral hypothalamus. (B) Acquisition performance before chemogenetic manipulation. (C) Responding to the previously reinforced stimulus during extinction, with MCH-DREADD activation restricted to this phase. (D) Sessions required to reach ≤30% previously reinforced responding. Control and MCH-DREADD, n = 8 mice/group. Acquisition and extinction trajectories were analyzed using mixed-effects models; extinction criterion was compared using Welch’s t-test. No group or group × session effect was detected during extinction. Data are mean ± SEM; individual animals are shown for the criterion measure.

Despite the robust effect of MCHR1 blockade during extinction, we detected no effect of LH-MCH neuronal activation on extinction behavior. Responding declined similarly in both groups across sessions (p < 0.001), with no group effect (p = 0.840) or group × session interaction (p = 0.418), and mice reached the extinction criterion after a comparable number of sessions (Control: 4.50 ± 0.27; MCH-DREADD: 5.13 ± 0.79; p = 0.47; Fig. 3). Acquisition before activation showed no group effect (p = 0.998) or group × session interaction (p = 0.239; Fig. 3B).

Thus, chemogenetically activating LH-MCH neurons during loss of reinforcement did not reproduce a reciprocal counterpart of the MCHR1-blockade phenotype. This dissociation suggested that receptor blockade and activation of the MCH neuronal population do not represent simple loss- and gain-of-function versions of the same process. We therefore asked whether neuronal activation becomes behaviorally relevant when contingency change requires active selection of a newly rewarded alternative.

### LH-MCH neuronal activation alters reversal learning and outcome-dependent choice

The absence of an extinction effect did not exclude a role for MCH-neuron activation during reversal, where successful adaptation requires active replacement of one rewarded stimulus by another. We therefore examined two successive reversals: a first reversal performed during chemogenetic activation of LH-MCH neurons, followed by a post-activation reversal (PAR) performed without MCH-DREADD activation (Fig. 4A). Acquisition was completed without chemogenetic manipulation. After acquisition criterion was reached, mice received a single CNO challenge while the original discrimination remained unchanged; this pre-reversal control revealed no significant group-specific change in first-choice performance or correction trials per error (Supplementary Fig. 2A,B). Both groups then received CNO before each session of the first reversal, producing hM3Dq-mediated activation only in MCH-DREADD mice. After criterion was reached, the stimulus–reward assignments were reversed again for PAR. During PAR, Control mice received CNO whereas MCH-DREADD mice received vehicle, such that MCH-DREADD activation was absent. Thus, Fig. 4 tests behavior during an activation-associated reversal and during a subsequent reversal after activation had ceased.

**Figure 4.**
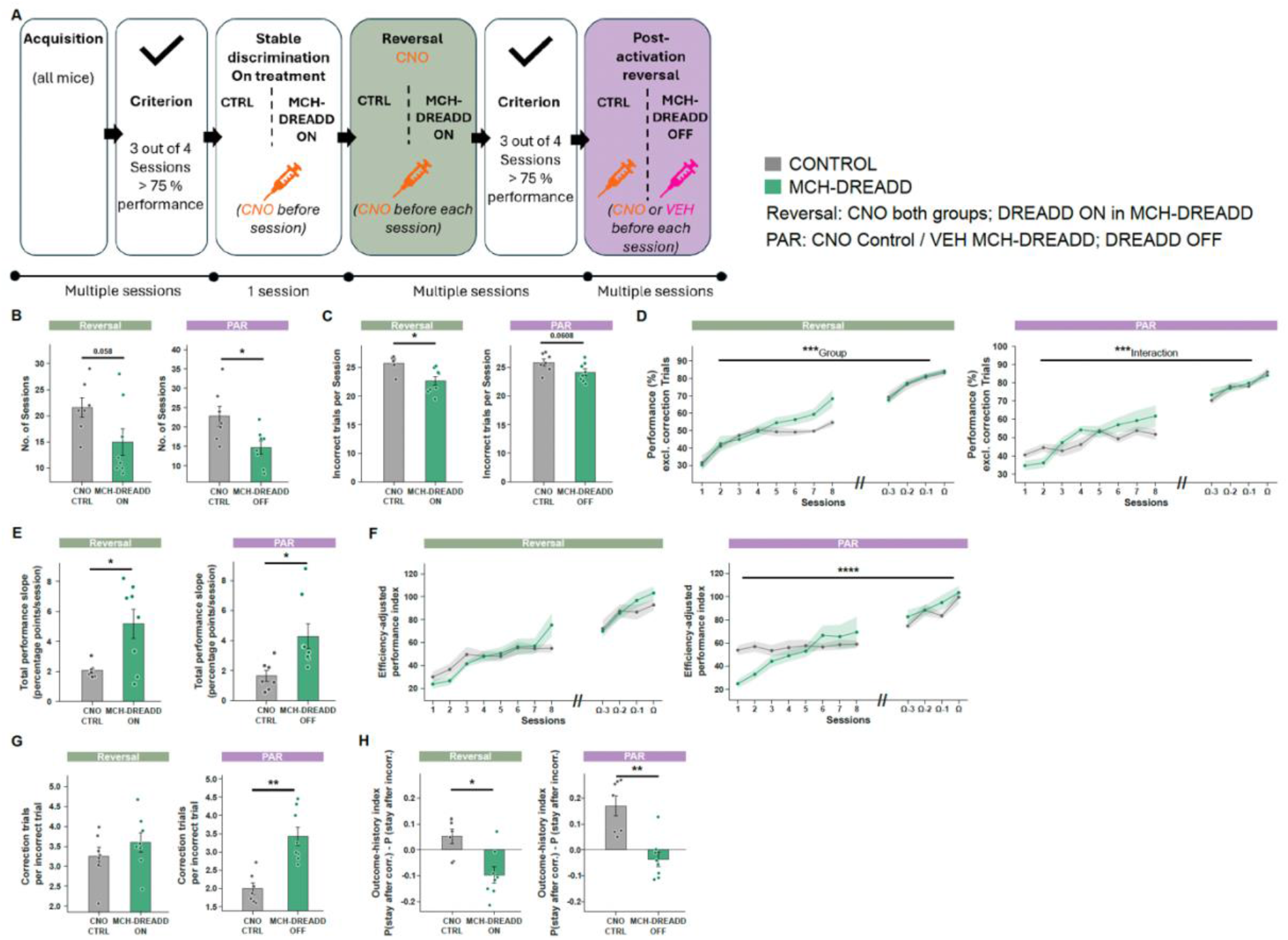
MCH-neuron activation alters reversal learning and outcome-dependent choice. (A) Experimental design. Acquisition was conducted without chemogenetic manipulation. The experiment comprised a first reversal performed during hM3Dq-mediated LH-MCH activation and a subsequent post-activation reversal (PAR) performed without MCH-DREADD activation. During the stable-discrimination challenge and the first reversal, both groups received CNO; hM3Dq-mediated LH-MCH neuronal activation occurred only in MCH-DREADD mice. During the subsequent post-activation reversal (PAR), Control mice received CNO and MCH-DREADD mice received vehicle; MCH-DREADD activation was therefore absent during PAR. (B) Sessions to criterion. (C) Incorrect regular trials per session. (D) First-choice performance excluding correction trials. (E) Per-animal total-performance slopes during the first reversal and PAR, estimated by linear regression of session-wise total performance within each phase. (F) Efficiency-adjusted performance index. (G) Correction trials per incorrect regular trial. (H) Outcome-history index, P(stay after correct) − P(stay after incorrect), where “stay” denotes repetition of the same touchscreen window choice on the subsequent trial. Control, n = 7; MCH-DREADD, n = 8. Longitudinal measures were analyzed by cluster-robust OLS with animal as cluster. Sessions-to-criterion comparisons were pre-specified and analyzed separately using two-sided Welch’s t-tests without cross-phase correction; remaining phase-specific summary comparisons were FDR-corrected. Data are mean ± SEM. *p < 0.05, **p < 0.01, ***p < 0.001, ****p < 0.0001.

During the first reversal, MCH-DREADD mice showed several indices consistent with faster acquisition of the new stimulus–reward relationship. MCH-DREADD mice tended to reach criterion in fewer sessions (Control: 21.57 ± 1.86; MCH-DREADD: 15.00 ± 2.54; p = 0.058; Fig. 4B), made fewer incorrect regular-trial choices (pFDR = 0.0165; Fig. 4C), performance overall (β = 7.66, p < 0.001; Fig. 4D). Per-animal total-performance slopes were also steeper (2.06 ± 0.18 versus 5.19 ± 0.98 percentage points/session; pFDR = 0.024; Fig. 4E). The efficiency-adjusted performance index showed neither a significant group effect (p = 0.133) nor group × session interaction (p = 0.302; Fig. 4F), indicating that neuronal activation did not uniformly improve every component of task performance.

During the second, post-activation reversal (PAR), group differences were evident across multiple measures of reversal learning. Although MCH neurons were no longer chemogenetically activated, animals previously exposed to activation reached criterion sooner (22.86 ± 2.60 versus 14.75 ± 1.67 sessions; p = 0.024; Fig. 4B) and showed higher first-choice performance with a steeper session-dependent increase (group: β = 8.72, p = 0.019; group × session: β = 1.23, p = 0.0005; Fig. 4D). Total-performance slopes were likewise higher (pFDR = 0.024; Fig. 4E), and the efficiency-adjusted performance index increased more steeply in the MCH-DREADD group (group × session: β = 2.13, p = 2 × 10⁻⁶; Fig. 4F). Thus, group differences remained evident after MCH-DREADD activation had ceased, although PAR cannot distinguish a persistent biological effect from a difference in learning history established during the first reversal.

We next asked whether this altered learning phenotype extended to the way individual outcomes controlled subsequent behavior. Correction trials per initial error were unchanged during reversal (pFDR = 0.457) but increased markedly in MCH-DREADD mice during PAR (Control: 1.99 ± 0.15; MCH-DREADD: 3.43 ± 0.25; pFDR = 0.00144; Fig. 4G). The outcome-history index, P(stay after correct) − P(stay after incorrect), where ‘stay’ denotes repeating the same touchscreen window choice on the subsequent trial, was also reduced in MCH-DREADD mice during both reversal (pFDR = 0.0128) and PAR (pFDR = 0.0093; Fig. 4H). Exploratory decomposition of this effect showed reduced win–stay behavior during reversal (pFDR = 0.033) and reduced lose–shift behavior during PAR (pFDR = 0.033; Supplementary Fig. 3A,B); the complementary comparisons did not survive FDR correction. The latter converged with the increased correction-trial burden during PAR, indicating greater persistence following negative feedback despite more rapid acquisition of the prevailing contingency.

Thus, MCH-neuron activation did not produce a uniform increase or decrease in behavioral flexibility. Instead, the pattern revealed a dissociation between contingency-level learning and the trial-by-trial influence of recent outcomes on subsequent choice.

## Discussion

The present study identifies the MCH system as a regulator of the balance between maintaining established reward contingencies and revising them when their consequences change. MCHR1 antagonism facilitated both appetitive extinction and reversal, with the clearest extinction effect occurring when receptor blockade coincided with contingency change. Chemogenetic activation of LH-MCH neurons produced a different pattern, with no detectable change in extinction but altered reversal learning and the influence of recent outcomes on subsequent choice. These findings distinguish MCHR1-mediated signaling from chemogenetic activation of the MCH neuronal population and argue against a simple model in which the MCH system acts along a single persistence– flexibility axis.

The timing experiment clarifies when MCHR1 signaling is most consequential. SNAP did not impair acquisition of the original appetitive response, whereas blockade during extinction produced a pronounced early reduction in responding once reward was withdrawn. The absence of a detectable SNAP effect during acquisition or during the stable-discrimination challenge also argues against a nonspecific reduction in reward value or task engagement as the primary explanation for its effects on updating. Together with our previous findings that MCHR1 antagonism facilitates extinction of active avoidance and reduces perseverative responding after a reward-guided rule switch (Kandasamey et al., 2026a,b), the present results point to a function that becomes particularly important when previously successful behavior is no longer appropriate. The convergence across aversive avoidance, appetitive extinction and reward reversal is notable because these tasks differ in valence and response requirements. What they share is the need to reduce the control exerted by an established contingency. The smaller effect of acquisition-only SNAP on the extinction criterion cautions against claiming absolute temporal exclusivity, but the trajectory data indicate that MCHR1 signaling is especially consequential during contingency revision itself.

The reversal data further define this effect. Reversal is not simply extinction with a new response added: the animal must suppress the previously rewarded choice while concurrently assigning value to its alternative, and contemporary accounts therefore regard reversal as the product of several interacting learning and control processes rather than a unitary measure of response inhibition (Izquierdo and Jentsch, 2012; den Ouden et al., 2013; Izquierdo et al., 2017; Soltani and Izquierdo, 2019). SNAP improved criterion attainment, the efficiency-adjusted performance index and the total-performance slope across reversal, without reducing correction trials per error. This pattern suggests that MCHR1 blockade facilitates adaptation to the reversed contingency without a detectable reduction in post-error perseveration, consistent with touchscreen work showing that overall reversal performance and post-error adjustment can dissociate (Marquardt et al., 2017; Dumont et al., 2025).

The chemogenetic experiments show that neuronal activation cannot be interpreted as the inverse of MCHR1 blockade. Activating LH-MCH neurons did not slow appetitive extinction, despite the strong effect of receptor antagonism. This is compatible with an increasingly heterogeneous picture of MCH-system function. Endogenous MCH-neuron activity contributes to object memorization and fear-extinction processes (Kosse and Burdakov, 2019; Concetti et al., 2020). Notably, Concetti et al. (2020) disrupted naturally occurring MCH-neuron activity associated with aversive events in a Pavlovian fear paradigm, whereas the present experiment increased population activity during appetitive instrumental extinction; differences in behavioral context, temporal precision and direction of manipulation therefore preclude a simple expectation of reciprocal effects. MCH neurons also participate in appetitive and consummatory processing (Noble et al., 2018; Terrill et al., 2020; Lord et al., 2021; Subramanian et al., 2023; Potter et al., 2026). Importantly, MCH neurons themselves engage more than MCH peptide signaling: most are glutamatergic, and glutamate and MCH can make partially dissociable contributions to their function (Schneeberger et al., 2018). Thus, chemogenetic activation engages a distributed cellular output that is mechanistically different from selectively blocking MCHR1. Consistent with a non-linear relationship between MCH-system manipulation and behavioral output, bidirectional perturbations of MCH/MCHR1 signaling can produce related changes in inhibitory control (Noble et al., 2019). These observations are consistent with the present conclusion that receptor- and cell-level manipulations need not yield reciprocal phenotypes.

The serial-reversal experiment further showed that group differences remained evident after MCH-neuron activation had ceased. MCH-DREADD mice showed evidence of faster acquisition of the changed contingency during the first reversal, and an even clearer phenotype emerged during PAR, after chemogenetic activation had ceased. This should not be interpreted as evidence that DREADD activation itself persisted. A more conservative possibility is that MCH-neuron activation during the first reversal altered the learning state with which animals encountered the next contingency change—for example, by changing the strength of the newly acquired contingency or by engaging learning-to-learn processes that emerge across serial reversals (Dickson et al., 2013; Kiyama et al., 2025). The accompanying choice-history results are informative in this respect. MCH-DREADD mice showed a reduced outcome-history index during both phases; separate analyses revealed reduced win–stay behavior during reversal and reduced lose–shift behavior during PAR. During PAR, the latter coincided with more correction trials following an initial error. Animals therefore acquired the prevailing contingency rapidly while becoming more persistent following individual negative outcomes. This apparent dissociation is important: contingency learning and trial-by-trial adjustment need not change in parallel. This distinction is consistent with touchscreen reversal work showing that recent outcome history and perseverative choice can be dissociated from overall reversal performance (Marquardt et al., 2017).

The translational relevance of these findings lies in the behavioral construct rather than in MCHR1 antagonism as a proposed treatment. Reversal-learning abnormalities and altered orbitofrontal-striatal recruitment have been reported in obsessive–compulsive disorder (Remijnse et al., 2006), and behavioral flexibility is among the domains with the strongest current cross-species comparability (Martins et al., 2025). By identifying MCH-system mechanisms that determine how reward history continues to control choice after contingencies change, the present study links hypothalamic function to a cognitive phenotype that can be interrogated in related forms across species.

Several boundaries remain. The pharmacological and chemogenetic experiments interrogate different levels of the MCH system and do not identify the circuit responsible for the touchscreen phenotype. Our previous active-avoidance work implicates prelimbic MCHR1 signaling in extinction, but the relevant receptor populations for appetitive updating remain unknown. Likewise, the PAR design establishes that behavioral differences are expressed after MCH-neuron activation has ceased, but cannot separate persistent biological consequences from the learning history created during the preceding reversal. What the present study establishes is that MCHR1 signaling favors the continued influence of established reward contingencies, whereas chemogenetic activation of LH-MCH neurons has broader, history-dependent effects on how contingencies and individual outcomes guide subsequent behavior. This distinction provides a framework for linking hypothalamic regulation of reward history to the control of behavioral persistence and adaptation across changing environments.

## Acknowledgments

We thank Scott E. Kanoski for providing the MCH-DREADD viral vector used in this study. This work was supported by ETH Zurich Grant ETH-24 20-2, awarded to D.P.-R.

## Data availability

Source data and analysis code will be deposited in a publicly available Open Science Framework (OSF) repository before publication.

---

**Supplementary Figure 1.**
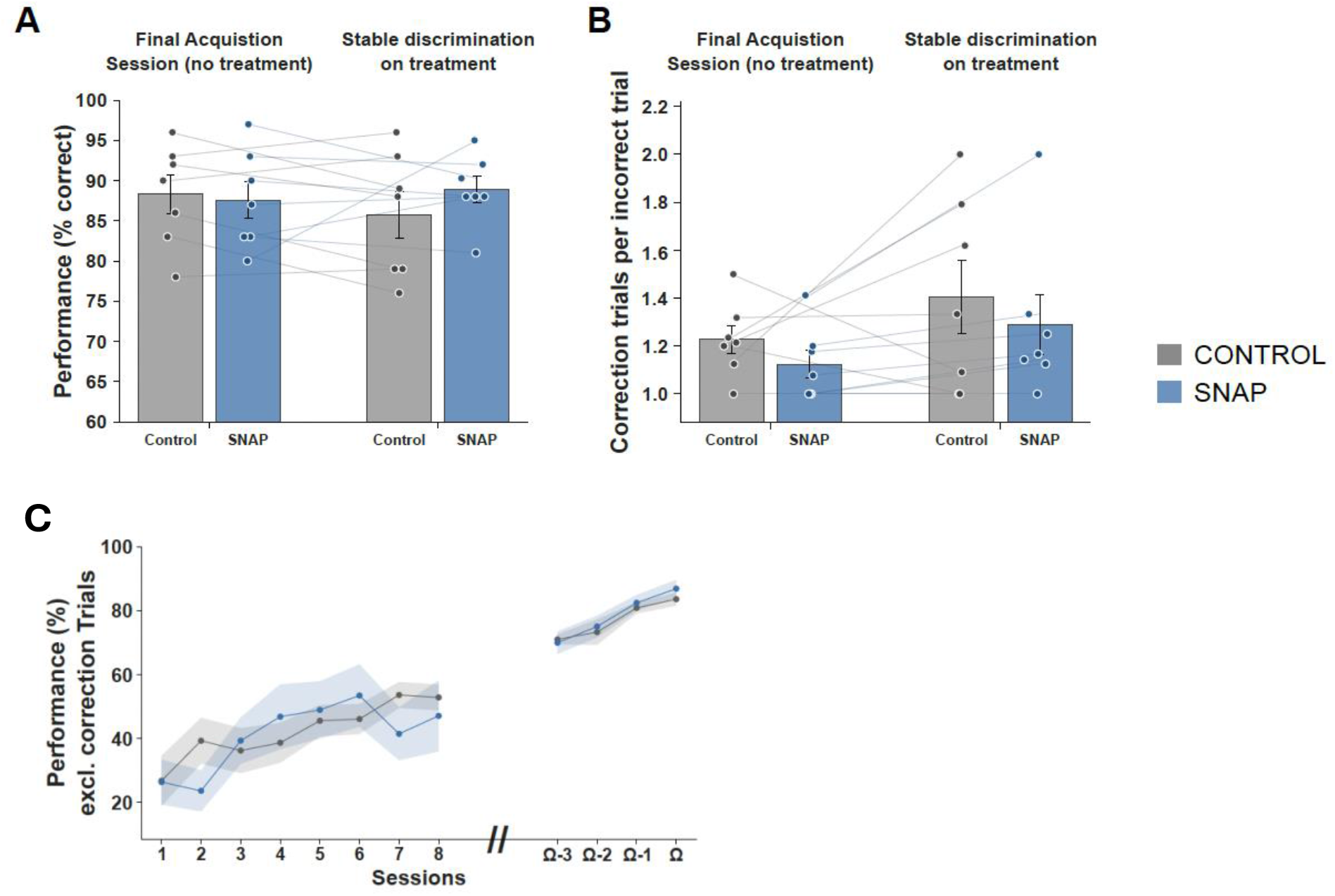
Additional performance measures in the SNAP visual-discrimination and reversal experiment. (A) First-choice performance during the final acquisition session and subsequent stable-discrimination treatment session. Accuracy was calculated from regular trials only, excluding correction trials. (B) Correction trials per incorrect regular trial during the same sessions. Within-group changes were tested using paired t-tests, and treatment-related change scores were compared between groups using Welch’s t-tests with FDR correction. No significant group-specific treatment effects were detected. (C) First-choice performance across reversal, calculated as the percentage of correct regular trials with correction trials excluded. Performance increased across sessions (β = 2.78, p < 0.001), but neither the overall group effect (β = 0.013, p = 0.998) nor the group × session interaction (β = 0.27, p = 0.790) was significant. Session-level data were analyzed by cluster-robust OLS with animal as cluster. Control and SNAP, n = 7 mice/group. Data are mean ± SEM.

**Supplementary Figure 2.**
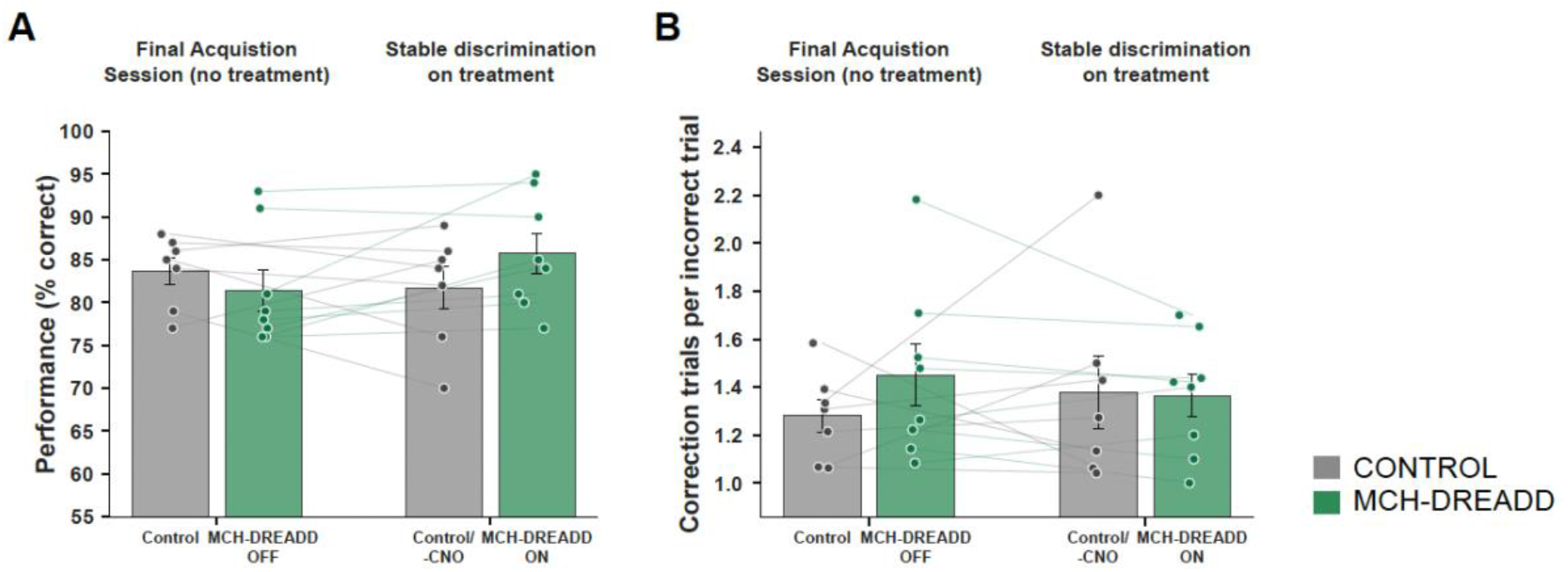
Acute LH-MCH neuronal activation under an established visual discrimination. (A) First-choice performance during the final acquisition session and the subsequent CNO treatment session. Accuracy was calculated from regular trials only, excluding correction trials. (B) Correction trials per incorrect regular trial during the same sessions. Control, n = 7; MCH-DREADD, n = 8. Both groups received CNO during the treatment session; hM3Dq-mediated MCH-neuron activation occurred only in MCH-DREADD mice. Lines connect repeated measurements from individual animals; bars show mean ± SEM. Within-group changes were tested using paired t-tests, and treatment-related change scores were compared between groups using Welch’s t-tests with FDR correction. Neither first-choice performance nor correction-trial burden showed a significant group-specific treatment effect.

**Supplementary Figure 3.**
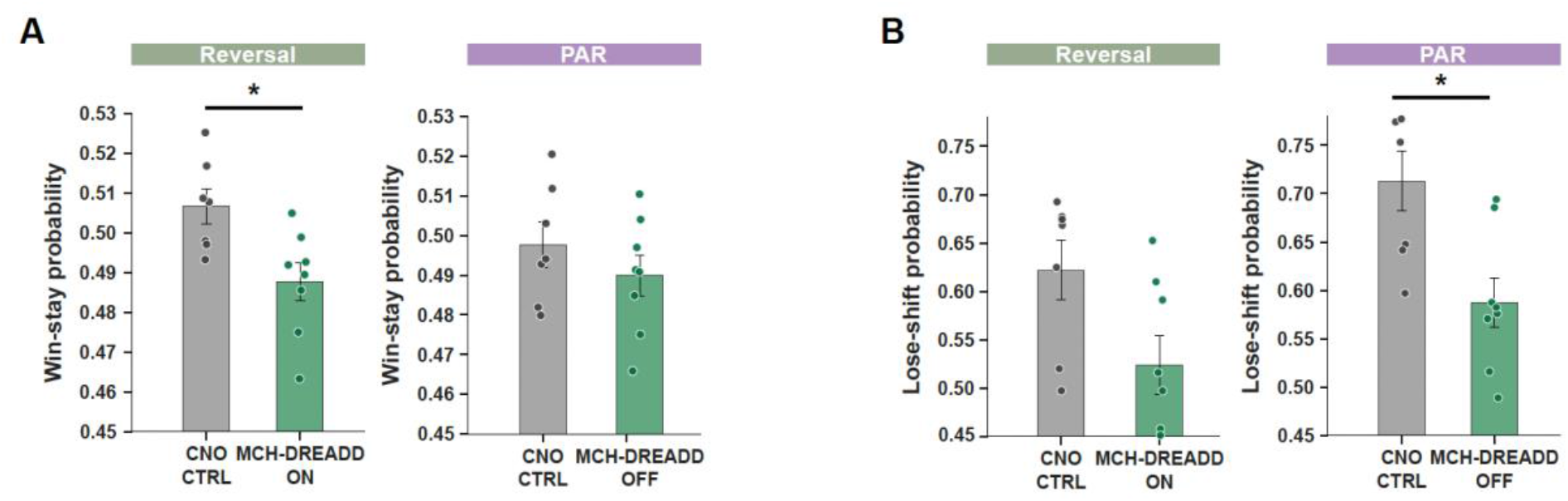
Outcome-dependent choice strategies during reversal and post-activation reversal. (A) Win–stay probability, defined as the probability of repeating the preceding window choice following a correct response. Win–stay probability was lower in MCH-DREADD mice during reversal (Control: 0.507 ± 0.004; MCH-DREADD: 0.488 ± 0.005; Welch’s t(13.00) = 2.96, pFDR = 0.033), but did not differ during PAR (Control: 0.498 ± 0.006; MCH-DREADD: 0.490 ± 0.005; t(12.64) = 1.01, pFDR = 0.360). (B) Lose–shift probability, defined as the probability of switching window choice following an incorrect response. The group difference during reversal did not survive FDR correction (Control: 0.622 ± 0.030; MCH-DREADD: 0.524 ± 0.030; t(12.91) = 2.30, pFDR = 0.078), whereas lose–shift probability was lower in MCH-DREADD mice during PAR (Control: 0.713 ± 0.031; MCH-DREADD: 0.588 ± 0.025; t(12.13) = 3.14, pFDR = 0.033). Probabilities were calculated per animal and session, including correction trials, and then averaged across sessions within each phase. LH-MCH neurons expressing hM3Dq were chemogenetically activated during reversal but not during PAR. Control, n = 7; MCH-DREADD, n = 8. Bars show mean ± SEM; points represent individual animals. Group comparisons used two-sided Welch’s t-tests with FDR correction.

## Materials and Methods

### Animals

Adult male C57BL/6J mice (≥8 weeks old at the start of behavioral testing) were used in all experiments. To maintain motivation for appetitive responding without conventional food or scheduled water restriction, mice had ad libitum access to 2% citric acid water, an established alternative to water restriction that reduces voluntary fluid intake while preserving motivation to obtain more palatable liquid rewards (Urai et al., 2021). Citric acid water was introduced 2 weeks before behavioral testing and remained available throughout the experiments. Standard food pellets were available ad libitum. Body weight was monitored regularly and maintained above 85% of pre-experimental body weight. Mice were housed on an inverted 12-h light/dark cycle, and behavioral testing was conducted during the dark phase. All procedures were performed in accordance with the requirements of the Zurich Cantonal Veterinary Office and the Animal Welfare Ordinance of the Swiss Federal Food Safety and Veterinary Office.

### Systemic pharmacology

For systemic MCHR1 antagonism, SNAP-94847 hydrochloride (Tocris Bioscience, Bio-Techne) was dissolved in vehicle containing 2% dimethyl sulfoxide (DMSO; PanReac AppliChem) and 25% β-cyclodextrin (Merck) in phosphate-buffered saline (PBS). SNAP-94847 was prepared at 2.67 μg/μL and administered intraperitoneally at 20 mg/kg.

For hM3Dq-mediated chemogenetic experiments, clozapine-N-oxide (CNO; Sigma-Aldrich) was prepared in 0.5% DMSO in saline and administered intraperitoneally at 0.5 mg/kg. Injection volume was calculated individually from body weight on the day of testing.

Whenever pharmacological or chemogenetic treatment was applied, injections were administered 45 min before the start of the behavioral session. Mice were weighed before each injection to calculate the appropriate injection volume.

### Stereotaxic viral injections

For chemogenetic experiments, anesthesia was induced with isoflurane and maintained at 1.5–2.0% throughout surgery.

Perioperative analgesia and topical lidocaine were administered to minimize pain. Mice were positioned in a stereotaxic frame, and the lateral hypothalamic region was targeted bilaterally. Viral injections were made using a 33-gauge needle mounted on a Hamilton syringe at the following coordinates relative to bregma: AP −1.35 mm, ML ±1.90 mm, DV −5.30 mm, with a 10° injection angle. The viral vector was AAV2-MCHp-hM3D(Gq)-mCherry-WPRE (Vector Biolabs; AAV2 capsid; titer, 7.4 × 10¹² genome copies/mL; lot 220207#42), in which hM3D(Gq)-mCherry expression was driven by the MCH promoter. A volume of 400 nL was infused per hemisphere at 50 nL/min. This MCH promoter-driven hM3Dq vector has previously been shown to selectively target MCH neurons and to increase their firing in response to CNO (Noble et al., 2018).

Control mice underwent a corresponding mock surgical procedure with the same anesthesia, perioperative analgesia, stereotaxic positioning, bilateral burr holes, and needle insertion at the lateral hypothalamic target coordinates, but without viral infusion.

Mice recovered for at least 3 weeks before behavioral testing to allow viral transduction and hM3Dq-mCherry expression.

### Histology

Following completion of the experiments, animals were deeply anaesthetised with pentobarbital and transcardially perfused with PBS (pH 7.4), followed by 4% paraformaldehyde (PFA) in PBS. Brains were collected, post-fixed overnight in 4% PFA at 4°C, cryoprotected overnight in 30% sucrose in PBS, frozen on dry ice, and coronally sectioned at 100 μm using a cryostat.

Free-floating sections were incubated overnight at 4°C with gentle agitation in rabbit anti-melanin-concentrating hormone antibody (MCH; 1:1,000; Phoenix Pharmaceuticals, H-070-47) diluted in antibody dilution buffer, comprising PBS containing 3% bovine serum albumin (BSA) and 0.1% Tween-20.

The following day, sections were washed three times for 5 min each in PBS containing 0.1% Tween-20 (PBST) and incubated for 1–2 h at room temperature, with gentle agitation in the dark, with Cy3-conjugated donkey anti-rabbit IgG (H+L) secondary antibody (1:500; Jackson ImmunoResearch, 711-165-152) diluted in antibody dilution buffer. Sections were then washed three times for 5 min each in PBST and once in PBS.

Images were acquired using a fluorescence microscope (Eclipse Ti2, Nikon) to verify viral expression and anatomical targeting.

### Touch screen apparatus

All behavioral experiments were conducted in triangular Bussey–Saksida mouse touchscreen chambers (model 80614; Campden Instruments). The touchscreen chamber configuration is shown in Supplementary Fig. 4. The chambers were controlled using ABET II Touch software. The front wall contained the touchscreen, and the opposite wall contained a food magazine through which strawberry milkshake (Emmi, Switzerland; 78 kcal/100 ml, 13 g carbohydrate, 3.5 g protein, and 1.1 g fat per 100 ml) was delivered. Unless otherwise specified, reward delivery consisted of 7 μL of strawberry milkshake. Task-specific masks were positioned in front of the screen to restrict responses to the relevant response windows. Chambers were cleaned with 70% ethanol between animals and sessions.

**Supplementary Figure 4.**
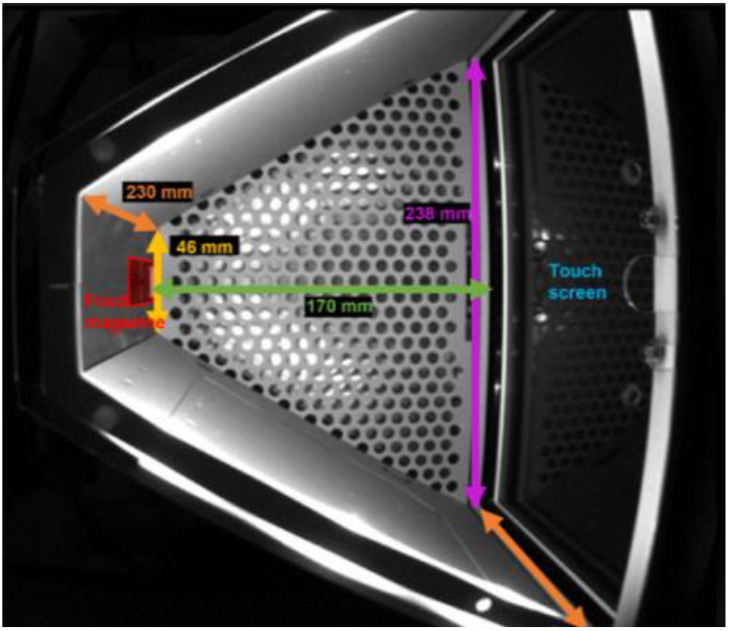
Touchscreen chamber. Triangular Bussey–Saksida mouse touchscreen chamber (Campden Instruments, model 80614) used for behavioral testing. The chamber contained a touchscreen at one end and a food magazine for delivery of strawberry milkshake at the opposite end. Task-specific masks restricted touchscreen responses to the appropriate response windows.

### Behavioral experiments

Before touchscreen training, mice were habituated to strawberry milkshake in the home cage.

Touchscreen responses and entries into the food magazine were detected automatically by infrared beam breaks. Mice completed no more than one behavioral session per day. Animals were randomly assigned to experimental groups before treatment. Behavioral responses and trial outcomes were recorded automatically by ABET II Touch software, and no manual behavioral scoring was used. Data analysis was performed with group identity masked until completion of the predefined analyses.

#### Extinction task

Mice were trained in a touchscreen-based extinction task consisting of acquisition followed by extinction. During acquisition, each session consisted of 30 trials. On each trial, a single white visual stimulus (4 x 4 cm) was presented on the touchscreen, and touching the stimulus was reinforced with 7 μL of strawberry milkshake paired with a 280-ms auditory cue and magazine-light illumination. Acquisition criterion was >95% stimulus-directed responding in at least 5 of 7 sessions. During extinction, the same visual stimulus was presented, but touchscreen responses to the stimulus were no longer reinforced. Mice completed 10 extinction sessions. Extinction was quantified as the decline in responding to the previously reinforced stimulus across sessions; the criterion measure was the number of sessions required to reach ≤30% previously reinforced responding.

To determine whether the behavioral effect of MCHR1 antagonism depended on treatment phase, mice were assigned to one of three groups. Control mice received vehicle before sessions during both acquisition and extinction. SNAP-Acq mice received SNAP-94847 during acquisition and vehicle during extinction, whereas SNAP-Ext mice received vehicle during acquisition and SNAP-94847 during extinction. Treatment was administered before each session of the corresponding phase.

In the chemogenetic extinction experiment, acquisition was completed without chemogenetic activation. During extinction, both MCH-DREADD and Control mice received CNO before each session; only MCH-DREADD mice expressed hM3Dq in MCH neurons and therefore underwent chemogenetic activation of the MCH neuronal population.

#### Visual discrimination (VD) task

The visual-discrimination/reversal procedure followed established touchscreen principles (Horner et al., 2013; Turner et al., 2017). The sequence of habituation, pretraining, acquisition and subsequent contingency manipulation is summarized in Supplementary Fig. 6. During all phases, a mask containing two response windows (7.5 × 7.5 cm; 0.5 cm apart) was positioned in front of the touchscreen. Mice were first habituated to the chamber for one 10-min session and to strawberry milkshake in the home cage. Food-magazine habituation was then conducted over three sessions (two 20-min sessions followed by one 30-min session). During magazine habituation, 7 μL of milkshake was delivered every 10 s and paired with illumination of the magazine and a 280-ms auditory cue. Entry into the magazine, detected by an infrared beam break, was scored as reward collection and terminated the magazine light.

##### Pretraining comprised four stages

Initial touch, Must touch, Must initiate, and Punish incorrect. Unless otherwise stated, reward delivery consisted of 7 μL of strawberry milkshake paired with a 280-ms tone and magazine-light onset.

###### Initial touch (30 min or 100 trials maximum per session; 3 sessions)

A white square stimulus was presented pseudorandomly in the left or right response window. Touching the stimulus, or 30 s without a response, terminated the stimulus and initiated reward delivery together with a 5-s intertrial interval (ITI). The magazine light was extinguished at the end of the ITI or when the reward was collected. During sessions 2 and 3, a stimulus touch was reinforced with 21 μL of milkshake, whereas 7 μL was delivered following a 30-s omission, to encourage interaction with the screen.

###### Must touch (30 min or 100 trials maximum per session; 4 sessions)

A white square was presented pseudorandomly in one of the two response windows. Reward was delivered only after the mouse touched the stimulus. Stimulus offset and reward delivery initiated a 5-s ITI, and the magazine light was extinguished at the end of the ITI or upon reward collection.

###### Must initiate (60 min or 100 trials maximum per session; 3 sessions)

At the start of each trial, the magazine light was illuminated. A magazine entry extinguished the light and initiated stimulus presentation. Touching the stimulus triggered stimulus offset, milkshake delivery, the auditory cue, and magazine-light onset. Reward collection extinguished the magazine light and initiated a 5-s ITI, after which the magazine light was illuminated to signal availability of the next trial.

###### Punish incorrect (60 min or 100 trials maximum per session; minimum 3 sessions; criterion >75% accuracy in 1 session)

Trial initiation and correct responses were reinforced as in the Must initiate stage. A response in the blank response window was scored as incorrect, terminated the stimulus, and initiated a 5-s timeout signaled by the house light, with no reward delivered. The same stimulus configuration was then repeated as a correction trial. Correction trials were repeated until a correct response was made and were not counted as regular trials.

###### Acquisition (60 min or 100 regular trials maximum per session; minimum 5 sessions; criterion >75% accuracy in 3 of 4 sessions)

Two brightness-matched visual stimuli (Supplementary Fig. 5) were presented within the two touchscreen response windows (7.5 × 7.5 cm; 0.5 cm apart). For each mouse, one stimulus was assigned as rewarded (S+) and the other as unrewarded (S−), with stimulus identity counterbalanced across animals. On each trial, the left/right position of S+ was pseudorandomized. Selection of S+ produced 7 μL of milkshake together with the 280-ms tone cue and magazine-light onset. Selection of S− initiated a 5-s timeout with house-light illumination and no reward, followed by a correction trial in which the same stimulus locations were repeated. Correction trials continued until S+ was selected and were excluded from regular-trial accuracy and the session criterion.

**Supplementary Figure 5.**
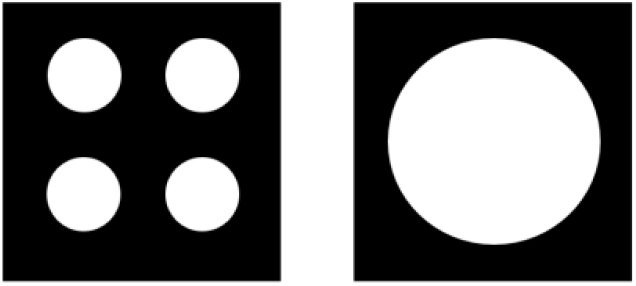
The two visual-discrimination stimuli. Visual stimuli used for the visual-discrimination and reversal task. One stimulus was assigned as rewarded (S+) and the other as unrewarded (S−), with stimulus identity counterbalanced across animals.

The S+/S− contingency remained unchanged throughout acquisition. Only mice that reached the prespecified acquisition criterion proceeded to the treatment challenge and reversal, ensuring a common performance threshold before pharmacological or chemogenetic manipulation. First-choice performance was calculated as correct regular trials divided by total regular trials × 100; correction trials were excluded. No injections were given during acquisition so that formation of the initial discrimination was not pharmacologically or chemogenetically manipulated. In the SNAP visual-discrimination cohort, two of 16 mice failed to reach acquisition criterion and were excluded from subsequent treatment-challenge and reversal analyses (one Control and one SNAP mouse), yielding *n* = 7 mice per group.

###### Stable discrimination on treatment (60 min or 100 trials maximum; 1 session)

After acquisition criterion was reached, mice completed one additional session under the unchanged discrimination contingency. In the SNAP experiment, mice received vehicle or SNAP-94847 according to group assignment. In the chemogenetic experiment, both MCH-DREADD and Control mice received CNO before this stable-discrimination challenge session; hM3Dq-mediated activation occurred only in MCH-DREADD mice. This session was used to determine whether the acute manipulation altered performance while the established stimulus–reward contingency remained valid.

###### Reversal (60 min or 100 trials maximum per session; criterion >75% accuracy in 3 of 4 sessions)

The task structure was identical to acquisition, but the stimulus–reward assignments were reversed: the former S+ became S−, and the former S− became S+. In the SNAP experiment, mice received vehicle or SNAP-94847 before each reversal session according to group assignment. In the chemogenetic experiment, both MCH-DREADD and control mice received CNO before each reversal session; hM3Dq mediated activation occurred only in MCH-DREADD mice. Correction trials were implemented as during acquisition and excluded from regular-trial accuracy.

Post-activation reversal (PAR; chemogenetic experiment only; 60 min or 100 trials maximum per session; criterion >75% accuracy in 3 of 4 sessions). After criterion was reached in the first reversal, the stimulus–reward contingencies were switched again. During PAR, Control mice received CNO whereas MCH-DREADD mice received vehicle; thus, hM3Dq-expressing MCH neurons were not chemogenetically activated during this phase. The stimulus rewarded during the first reversal became unrewarded, and the alternative stimulus became rewarded.

**Supplementary Figure 6.**
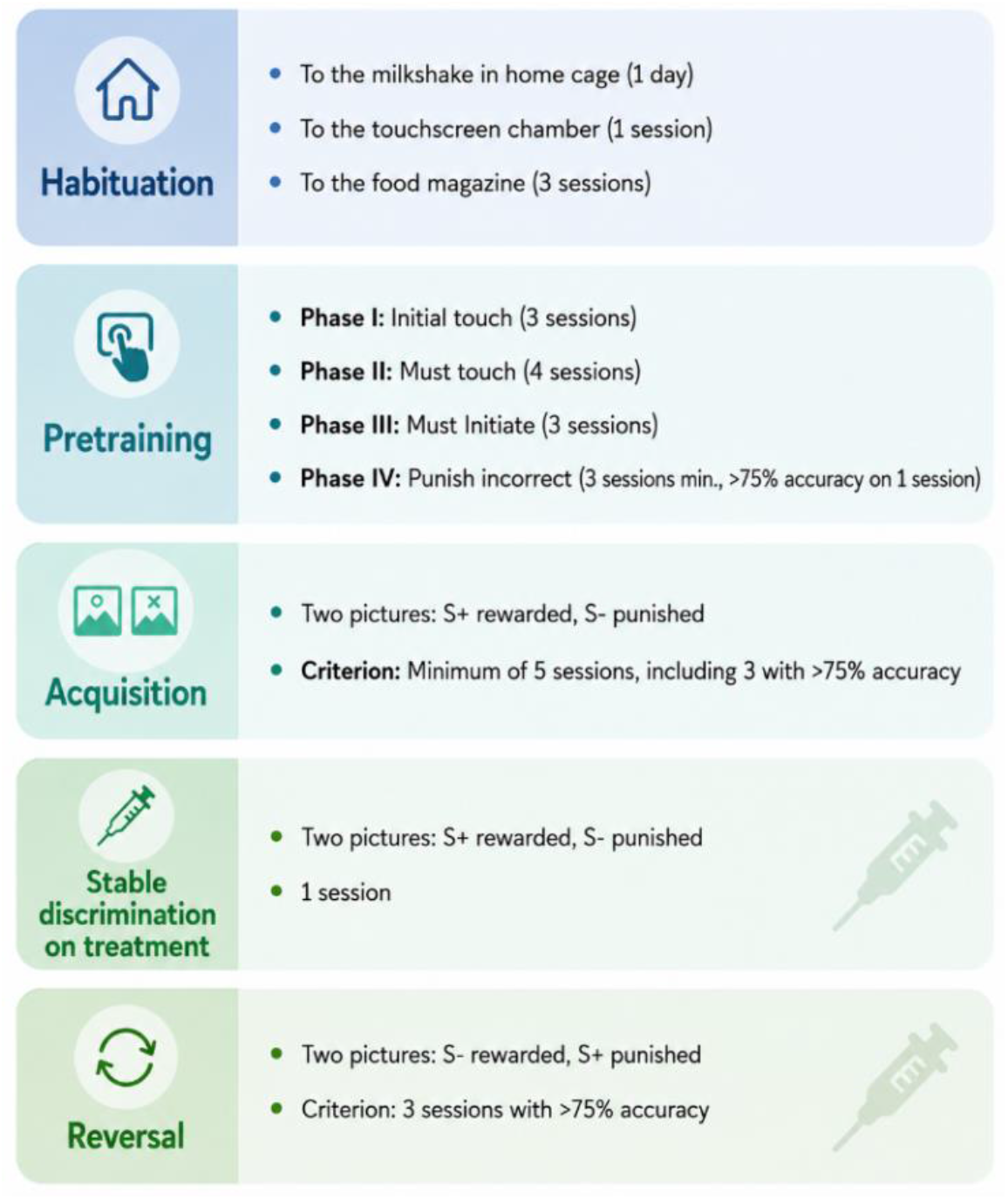
Touchscreen training/task sequence. Behavioral training sequence for the visual-discrimination/reversal experiments, comprising habituation, touchscreen pretraining, visual-discrimination acquisition, a stable-discrimination treatment challenge, and reversal. Following acquisition to criterion, pharmacological or chemogenetic manipulations were introduced according to experimental group and task phase.

### Statistical analysis

Statistical analyses and graphical visualizations were performed in Python 3.9. Data are reported as mean ± SEM unless otherwise stated, and statistical significance was set at p < 0.05. The mouse was the experimental unit. Analyses of session-level data accounted for repeated observations within animal. Exact sample sizes and statistical tests are reported in the corresponding figure legends.

#### Extinction analyses

Acquisition and extinction trajectories were analyzed using session-level mixed-effects models including treatment group, session, and their interaction. For the SNAP timing experiment, an additional model spanning acquisition and extinction included behavioral phase to determine whether treatment-dependent changes were specific to the transition from acquisition to extinction. Sessions required to reach the extinction criterion were compared using Welch’s t-tests. In the SNAP experiment, the two comparisons of SNAP-Acq and SNAP-Ext with the common Control group were corrected for multiple testing using the Benjamini–Hochberg false-discovery-rate procedure (Benjamini and Hochberg, 1995). For the MCH-DREADD extinction experiment, the single Control versus MCH-DREADD criterion comparison was analyzed using a two-sided Welch’s t-test.

#### Visual-discrimination and reversal analyses

First-choice performance was defined as the percentage of correct regular trials, with correction trials excluded. Correction-trial burden was calculated as the number of correction trials relative to incorrect regular trials. For the stable-discrimination treatment challenge, changes from the final acquisition session to the treatment session were assessed within groups using paired t-tests, and treatment-related change scores were compared between groups using Welch’s t-tests. Benjamini–Hochberg FDR correction was applied to the predefined challenge comparisons.

Sessions required to reach reversal criterion in the SNAP experiment were compared using a pre-specified one-tailed Welch’s t-test based on the directional hypothesis that MCHR1 antagonism would facilitate contingency updating. Session-wise first-choice performance and the efficiency-adjusted performance index were analyzed using ordinary least-squares models with cluster-robust standard errors clustered by animal. Models included group, session, and the group × session interaction. Total performance was calculated as (correct regular trials + correct correction trials)/(regular trials + correction trials) × 100. The efficiency-adjusted performance index was obtained by scaling total performance by the median total trial count for the corresponding phase/session divided by the animal’s total trial count. Individual total-performance slopes were estimated for each animal by fitting a linear regression to session-wise total performance, including regular and correction trials. In the SNAP reversal experiment, total-performance slopes were compared between groups using a two-sided Welch’s t-test.

In the MCH-DREADD experiment, sessions-to-criterion comparisons for reversal and PAR were pre-specified phase-specific outcomes and were analyzed separately using two-sided Welch’s t-tests without cross-phase correction. Incorrect regular trials were averaged across sessions within each phase. Correction-trial burden was calculated for each animal and phase as the total number of correction trials divided by the total number of incorrect regular trials. Individual total-performance slopes were compared between groups using two-sided Welch’s t-tests. For each behavioral outcome, group comparisons were performed separately within each phase, and Benjamini–Hochberg FDR correction was applied across all phase-specific comparisons included for that outcome. Thus, each behavioral outcome constituted a separate FDR family; the number of comparisons entering the correction depended on whether that outcome was analyzed across acquisition, reversal and PAR or only across reversal and PAR. Adjusted values are reported as pFDR.

The outcome-history index was calculated as P(stay after correct) − P(stay after incorrect), with positive values indicating greater choice persistence following a correct than an incorrect trial. Win–stay and lose–shift probabilities were analyzed separately as exploratory decompositions of this index. For each strategy measure, phase-specific Control versus MCH-DREADD comparisons were performed using two-sided Welch’s t-tests and corrected using the same Benjamini–Hochberg FDR procedure.

